# Inverse covariation between the neocortex and the limbic system is concentrated at deep evolutionary divergences

**DOI:** 10.64898/2026.10.19.760202

**Authors:** Benjamin Wu, Nabil Imam

## Abstract

The macroscale architecture of mammalian brains consists of a broadly conserved set of structures, but the relative sizes of these structures vary substantially across species. Much of this variation follows conserved scaling relationships, but particular systems can expand or contract relative to the rest of the brain, providing targets for adaptive selection. One prominent example is a volumetric tradeoff between the neocortex and the limbic system, in which expansion of one is associated with reduction of the other, a feature apparent when brain structure volumes are examined across living species. However, whether this relationship has operated throughout mammalian evolution or arose during particular periods of diversification is unknown. Using phylogenetic comparative methods, we here show that the tradeoff between the neocortex and the limbic system is concentrated at deep evolutionary divergences separating major mammalian groups. After accounting for shared ancestry and brain size, the tradeoff strengthens progressively among deeper branches of the mammalian phylogeny, extending broadly across limbic structures, with the strongest signal involving olfactory components. Recent divergences show no consistent relationship. Our analyses indicate that the neocortex-limbic tradeoff emerged during the early diversification of mammals, highlighting how present-day variation in brain organization retains signatures of ancient evolutionary transitions.

---

Across mammals, brain structures vary widely in size, but this variation is not unconstrained. Major brain structures scale together in a manner consistent with shared developmental and genetic programs, a pattern often described as concerted evolution [1–3]. Despite this broad coordination, the sizes of different brain structures relative to one another can vary with the functional demands of a species [4, 5]. Such variation allows natural selection to emphasize some functions over others without enlarging the brain as a whole. A prominent example of this, at the level of major brain structures, is an inverse relationship between the neocortex and the limbic system, where volumetric expansion of one is associated with reduction of the other, reflecting a tradeoff in the allocation of neural tissue between two functionally and developmentally distinct systems [6, 7].

When during mammalian evolution this tradeoff arose, however, is unclear. Evidence for the neocortex-limbic covariation has come primarily from comparisons of brain structure volumes among living species without an examination of its possible origins in evolutionary history. This history encompasses divergences of vastly different depths [8–10], from recent splits between closely related species to ancient divergences separating major taxonomic groups. Distinguishing among these phylogenetic depths is necessary for determining whether the tradeoff represents a continuously operating feature of brain evolution or a transition concentrated at particular points in history.

Here we analyze the evolutionary relationship between the neocortex and the limbic system across 157 mammalian species spanning eight taxonomic groups. Using a time-calibrated evolutionary tree and phylogenetic comparative methods, we find that the apparent tradeoff between the two structures largely disappears when evolutionary divergences across the tree are pooled. Strikingly, the relationship re-emerges at deep evolutionary divergences and becomes progressively stronger with phylogenetic depth, reaching its greatest magnitude among the ancient branches separating major mammalian groups. At these deep divergences, the tradeoff extends across the entire olfactory-limbic complex, including the olfactory bulb, olfactory cortex, amygdala, septum, hippocampus, and parahippocampal regions. By contrast, recent divergences show no consistent tradeoff, nor do analyses within individual taxonomic groups. Our results indicate that the tradeoff between the neocortex and the limbic system is not a continuously operating constraint on mammalian brain evolution, but rather a signature of the reorganizational events that accompanied the earliest divergences among mammalian lineages.

## Volumetric covariation between brain structures

We compiled brain volume measurements of major brain structures from 157 mammalian species spanning eight taxonomic groups [6, 11–14]. We examined the relationship between each structure and a conserved set of core brain components comprising the medulla, mesencephalon, diencephalon, and striatum [15]. Among the structures in our dataset, we summed the volumes of the olfactory bulb, olfactory cortex, amygdala, septum, hippocampus, and schizocortex (which largely corresponds to the parahippocampal region, see Methods) for a combined volume of the limbic system [6, 7]. As expected under concerted scaling [1, 3, 15], the volumes of both the limbic system and the neocortex increase predictably with the volume of the core brain components (Figure 1A). The regression lines in Figure 1A define the allometrically expected volume of each structure given brain-core size, providing a baseline against which deviations from concerted scaling can be assessed.

**Figure 1:**
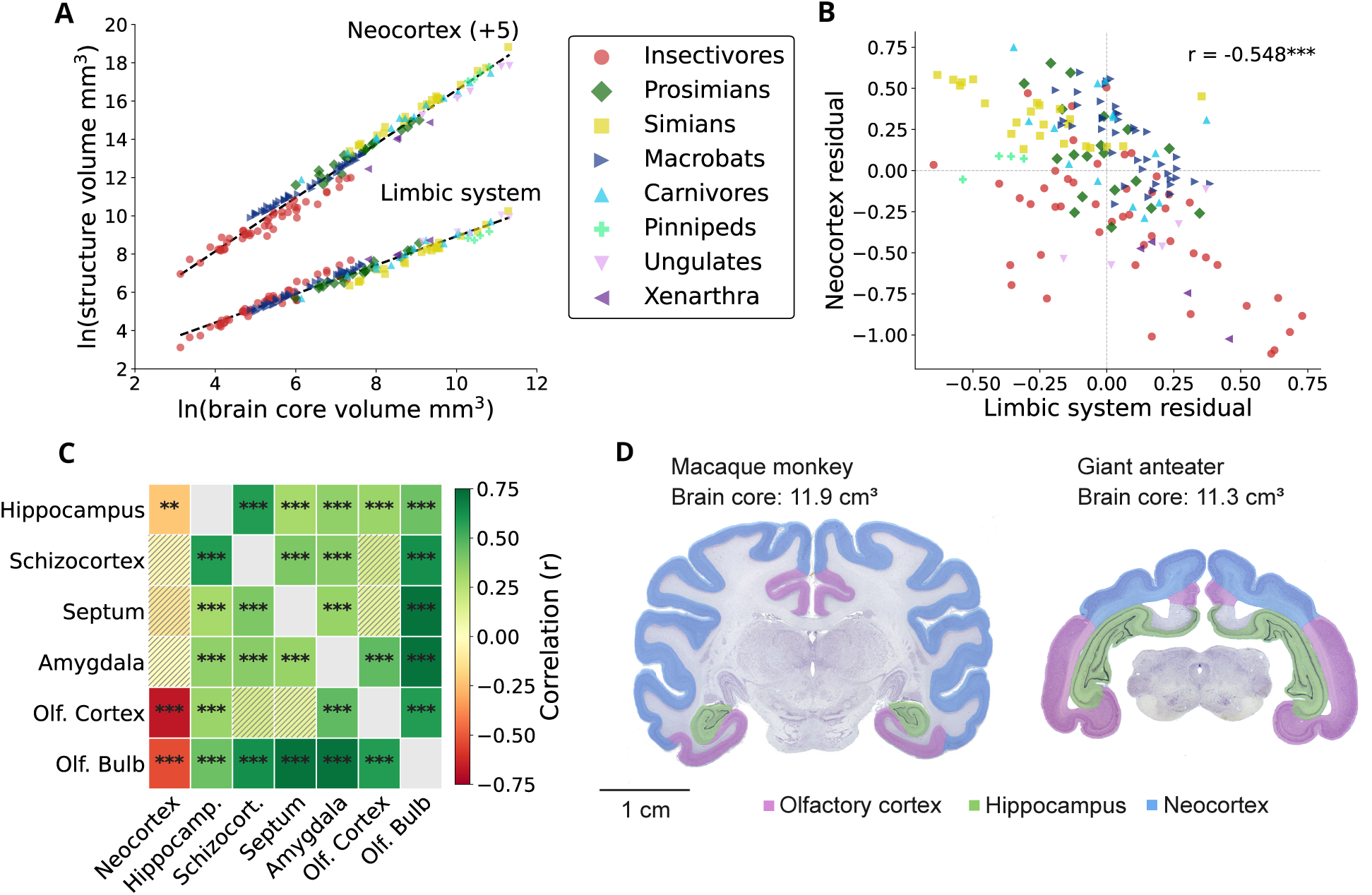
Volumetric covariation between major brain components. **(A)** Log-transformed neocortex and limbic system volumes (y-axis) plotted against brain-core volume (x-axis) across 157 mammalian species. The neocortex volumes are shifted by a constant (+5) on the *y* axis to separate the plots visually. As brain core expands, both the neocortex and the limbic system scale predictably. The regression line running through each set of scatter points provides the allometrically expected size of each structure, given brain-core size. **(B)** Residuals of the neocortex (y-axis) and the limbic system (x-axis) around their regression lines in subplot *A*, revealing a negative relationship between the two structures (r = -0.548, p *<* 0.001). **(C)** The volume of each limbic component and the neocortex were regressed individually against the volume of brain core and their residuals were computed. Shown are the pairwise correlations among those residuals. Individual limbic components covary positively with one another and negatively with the neocortex. **(D)** Coronal sections of the brains of a macaque monkey (Macaca mulatta) and a giant anteater (Myrmecophaga tridactyla), chosen to demonstrate contrasting allocation between neocortex and olfactory–limbic structures in two species with closely matched brain-core volumes. The coronal sections are from the Comparative Mammalian Brain Collections, www.brainmuseum.org, property of the University of Wisconsin and Michigan State Comparative Mammalian Brain Collections, funded by the National Science Foundation and the National Institutes of Health.

To quantify these deviations, we calculated residuals around the regression line of each structure. These residuals measure the relative enlargement or reduction of each structure compared with its expected volume. Examining these values, we observe that species with relatively enlarged limbic systems possess relatively smaller neocortices, and vice versa, producing a pronounced negative relationship (r=-0.548, p*<*0.001; Figure 1B), consistent with previous observations [6, 7]. This signal was driven primarily by the olfactory bulb, olfactory cortex, and the hippocampus, each of which individually covaried negatively with the neocortex (Figure 1C). In contrast, all of the individual limbic structures positively covaried with one another (Figure 1C), indicating that species with enlargement of one limbic component generally exhibit enlargement across all limbic structures. These patterns suggest that the limbic system and the neocortex participate in a coordinated evolutionary tradeoff, motivating a phylogenetically informed analyses, presented in the next section.

We next assessed whether these patterns hold within taxonomic groups by repeating the analysis within each taxon. The inverse relationship between the limbic system and the neocortex was not statistically significant in any individual taxon. Only insectivores showed a significant within-group tradeoff between the neocortex and one of the limbic structures—the olfactory cortex (*r* = −0.312, *p <* 0.05, *n* = 47). Covariation among limbic structures was similarly inconsistent within taxa. Only in insectivores did the septum and the amygdala each covary positively and significantly with all other limbic structures (Supplementary Figure 1). The signal of correlated evolution shown in Figure 1C therefore exists primarily between taxonomic groups, rather than within them, raising the possibility that the pattern arises from shared ancestry within the groups, instead of a tradeoff that operated repeatedly as lineages diverged.

## Disentangling shared ancestry from functional covariation

Brain volumes across species cannot be treated as independent observations because closely related species share common ancestry and therefore tend to resemble one another, potentially creating correlations between traits that do not reflect independent evolutionary change. For example, if two closely related species both possess larger-than-expected hippocampi and olfactory cortices, this does not necessarily imply coordinated expansion of the two structures. Rather, both species may simply have inherited the enlarged structures from a common ancestor. The apparent relationship between the structures could therefore reflect shared ancestry rather than repeated evolutionary covariation.

To determine whether the inverse relationship between the limbic system and the neocortex reflects shared ancestry or independent evolutionary change, we computed phylogenetic independent contrasts (PICs) across the evolutionary tree of the 157 species in our dataset (Figure 2A). PICs transform species trait values into contrasts at internal nodes of an evolutionary tree, where each contrast represents an independent divergence event [16]. We obtained a time-calibrated tree from the TimeTree database [10], which synthesizes published molecular timetrees and divergence-time estimates into an integrated evolutionary tree. Starting from log-transformed brain volumes at the leaves of the tree (extant species), we calculated contrasts for brain core, limbic system, and neocortex at each internal node using a post-order traversal of the tree (Methods). We then regressed the contrasts of each structure against the contrasts of brain core (Supplementary Figure 2). The residuals from these regressions represent brain structure volumes that account for both allometric scaling and phylogenetic non-independence.

**Figure 2:**
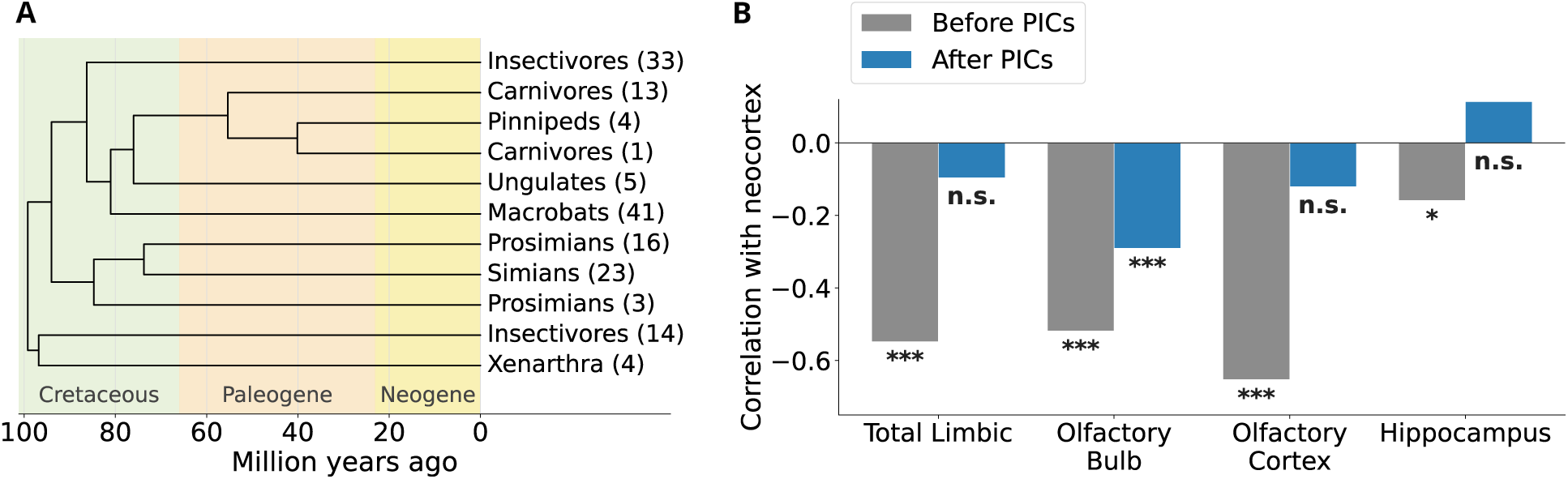
Phylogenetic corrections to the neocortex-limbic covariation. (A) Time-calibrated phylogeny of the major mammalian clades in our dataset. The number of species in each clade is shown in brackets (total 157 species). Insectivores and prosimians each appear in two separate branches because these groups are not monophyletic in the tree. Carnivores also appear twice because pinnipeds fall within the carnivore lineage but are shown separately. The full tree is shown in Supplementary Figure 4. (B) Bar plots of residual correlations between the neocortex and the limbic system before and after phylogenetic correction. The correlation is negative and statistically significant before correction (*r* = *−*0.548, *p <* 0.001), but weakens and loses significance afterward (*r* = *−*0.096, n.s.). Correlations between the neocortex and three individual limbic components—the hippocampus, olfactory cortex, and olfactory bulb—are also shown. These are the components that exhibited significant negative correlation with the neocortex before phylogenetic correction. After correction, only the olfactory bulb retains significance.

The PICs analysis shows that the inverse relationship between the limbic system and the neocortex described in the previous section weakens (r = -0.548 before PICs, r = -0.096 after PICs) and loses statistical significance after phylogenetic correction. Repeating the analysis for individual limbic components, we find that only the olfactory bulb maintains a significant relationship with the neocortex, although its negative correlation is substantially weakened (Figure 2B). The negative correlation between the olfactory cortex and the neocortex, which is strong and significant in the raw species data, becomes weak and nonsignificant after PICs. The relationship between the hippocampus and the neocortex also loses significance. In addition, the strong positive covariation among individual limbic structures in the raw species data weakens after phylogenetic correction, with some pairs losing statistical significance (Supplementary Figure 3). Together, these results indicate that much of the covariation observed in the raw species data reflects shared ancestry rather than independent evolutionary change. The olfactory bulb stands out as the one component for which an evolutionary relationship with the neocortex persists after phylogenetic correction.

## Deep evolutionary divergences reveal the neocortex-limbic tradeoff

The previous section pooled contrasts across all nodes of the evolutionary tree. Here we ask whether a stronger signal of correlated evolution is concentrated at particular depths of the tree.

We first performed a binary deep-versus-shallow comparison with a cutoff at 50 million years ago (Mya), partitioning divergences across the tree into a deep subset older than 50 Mya and a shallow subset more recent than 50 Mya, and analyzing brain-structure covariation within each (Figure 3). The deep subset contained 20 contrasts, whereas the shallow subset contained 136 contrasts. Analyzing these contrasts, we found that the tradeoff between the neocortex and the limbic system was concentrated among the deep divergences (Figure 3), where the two structures exhibited a pronounced negative relationship (*r* = −0.751, *p <* 0.001). In comparison, the relationship was weak and non-significant in the shallow subset (*r* = +0.072, n.s.). This partition was robust across thresholds of 40, 50, and 60 Mya (Supplementary Figure 5).

**Figure 3:**
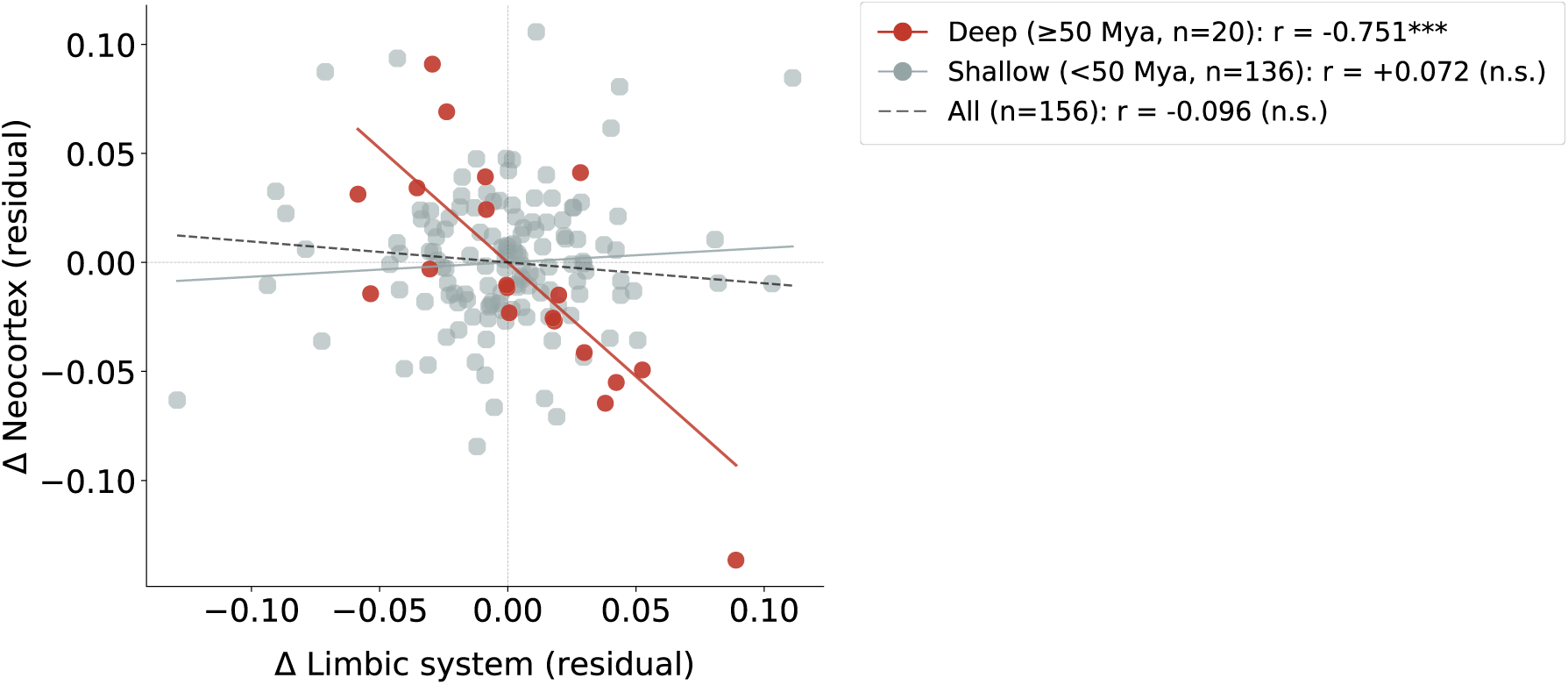
The tradeoff between the neocortex and the limbic system is concentrated at deep evolutionary divergences. Each point represents one divergence in the evolutionary tree and shows the evolutionary change in limbic system size (horizontal axis) and neocortex size (vertical axis) across that divergence, after removing the change expected from brain-core size. These size-corrected changes are the residuals of each structure’s contrasts after regression on brain-core contrasts. Divergences are partitioned into deep (*≥* 50 Mya, red, *n* = 20) and shallow (*<* 50 Mya, gray, *n* = 136) subsets. The inverse covariation between the two structures is strong at deep divergences (*r* = *−*0.751, *p <* 0.001), but absent at shallow ones (*r* = +0.072, n.s.).

Examining individual limbic components among the deep subset revealed a broadly shared pattern of negative covariation with the neocortex (Supplementary Figure 6). The olfactory bulb (*r* = −0.752, *p <* 0.01, *n* = 12 contrasts) and olfactory cortex (*r* = −0.735, *p <* 0.001, *n* = 20) showed the strongest negative relationships, followed by the septum (*r* = −0.576, *p <* 0.01, *n* = 20). The schizocortex (*r* = −0.431, *p* = 0.058, *n* = 20) and hippocampus (*r* = −0.411, *p* = 0.072, *n* = 20) exhibited weaker negative correlations that approached statistical significance. The amygdala showed a comparable negative correlation (*r* = −0.415, *p* = 0.18, *n* = 12), but did not reach significance, likely due to the smaller number of contrasts available to detect an effect of that size. Thus the tradeoff at deep nodes was not restricted to olfactory structures but extended broadly across the limbic system, with every component showing negative covariation with the neocortex, and the olfactory structures and septum reaching significance at *p <* 0.05. In the shallow subset, no consistent pattern was observed. The olfactory bulb retained a weak negative relationship (*r* = −0.183, *p <* 0.05), while the hippocampus covaried positively (*r* = +0.213, *p <* 0.05).

To assess how the strength of this covariation changed continuously with phylogenetic depth, we computed the correlation between the neocortex and the limbic system across a sweep of minimum node-age thresholds, beginning at 5 Mya and increasing in 2-million-year increments, including at each threshold only contrasts whose node ages exceeded the threshold. For example, at a threshold of 25 million years, the correlation was computed using only contrasts arising from nodes older than 25 Mya. We found that the negative correlation between the neocortex and the limbic system strengthened progressively with increasing threshold (phylogenetic depth) and then plateaued to approximately -0.75 (Figure 4A). We repeated the analysis for individual limbic components, and found a similar pattern, with correlations stabilizing to 0.74 for olfactory cortex, 0.76 for olfactory bulb, 0.58 for septum, 0.43 for schizocortex and 0.41 for hippocampus, at phylogenetic depths of approximately 48 Mya or deeper (Supplementary Figure 7).

**Figure 4:**
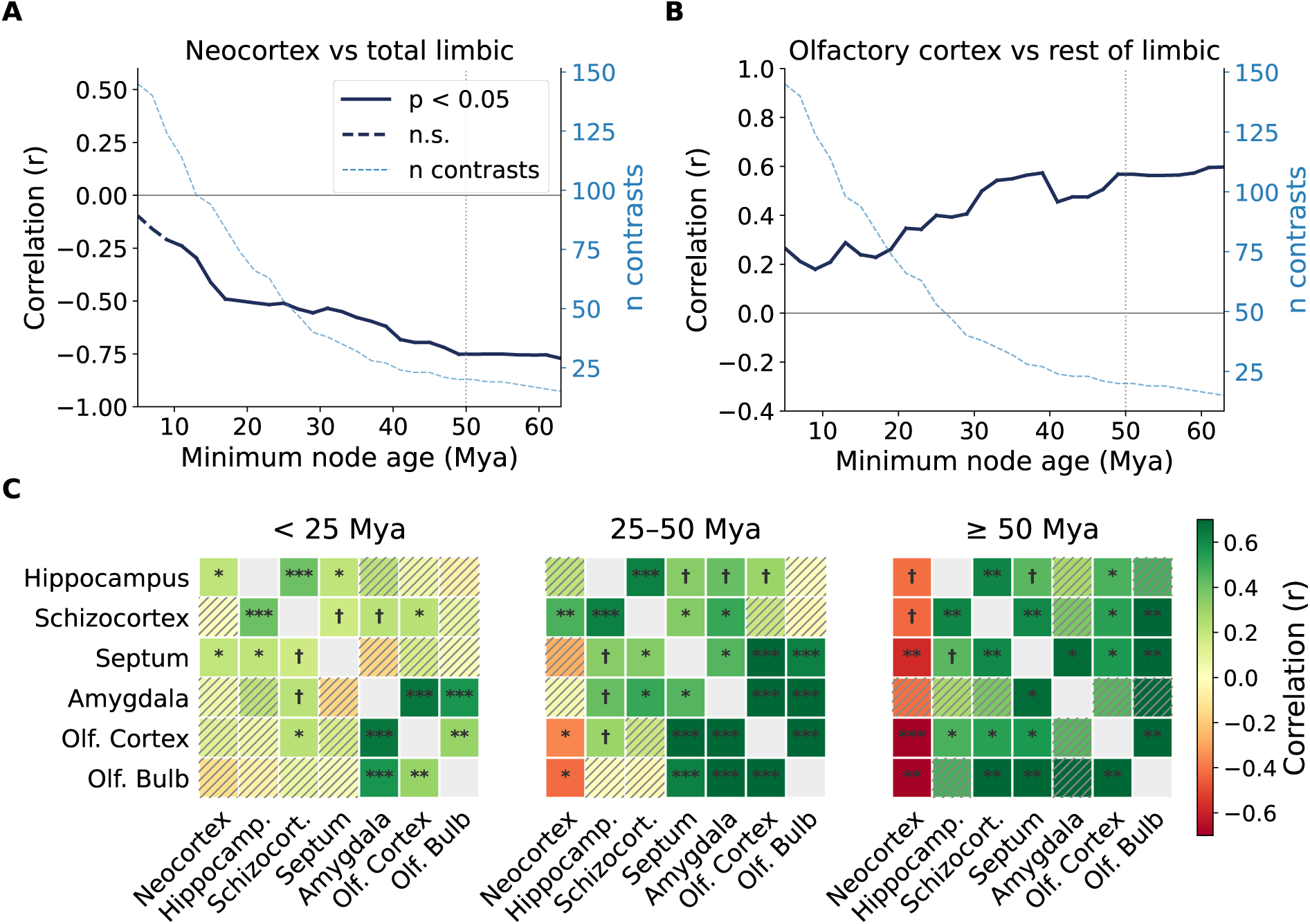
Neocortex–limbic and within-limbic covariation across evolutionary depth. (A) Residual correlation between the contrasts of the neocortex and the contrasts of the limbic system as a function of minimum node-age threshold. The correlation strengthens progressively with depth and plateaus approximately 48 Mya and earlier. Solid segments indicate *p <* 0.05, dashed segments indicate non-significance (n.s.). The number of contrasts at each threshold is shown (dashed blue line, right axis). (B) As in *A*, but for the olfactory cortex against the remaining limbic structures (olfactory bulb and amygdala are excluded, see main text). Unlike in *A*, the correlation here is positive and strengthens with depth. The correlation is strongest at the same divergences where the limbic structures trade off against the neocortex. (C) Pairwise residual correlations between the contrasts of each structure, partitioned by phylogenetic depth. Green denotes positive correlation, red negative, and stripes denote non-significance (*p ≥* 0.05). Correlations among limbic structures are positive in all three bins and generally increase in magnitude with depth. Correlations between the neocortex and limbic structures instead change sign, from positive or null among shallow divergences to uniformly negative at *≥* 50 Mya. Asterisks and daggers denote significance levels (see Methods).

We then examined how individual limbic structures covaried with one another at increasing depths. For this analysis, we used the olfactory cortex as a reference and computed its correlation with the combined volume of the septum, hippocampus, and schizocortex (the olfactory bulb and amygdala were excluded because of the fewer number of contrasts available for these structures). We found that this covariation strengthened progressively with depth, remaining positive and significant across thresholds (Figure 4B). The correlation stabilized at 0.57 at a depth of approximately 50 Mya or deeper.

To examine all pairs of structures, we next grouped contrasts into three depth bins (*<* 25 Mya, 25–50 Mya, and ≥ 50 Mya) and computed the correlation between every pair within each bin (Figure 4C). We found that correlations among limbic structures became stronger and more uniformly positive with increasing depth. In comparison, correlations between the neocortex and each limbic structure became uniformly negative at ≥ 50 Mya. Because the deepest bin contains fewer contrasts (Supplementary Figure 8), a few of the correlations, mostly involving the amygdala, did not reach statistical significance despite their magnitude. Although the amygdala’s deep correlations were noticeable, they were calculated using only 12 contrasts (4 for the amygdala–olfactory bulb pair), and therefore did not have enough statistical power to reach significance. Among the other structures, most deep correlations were strong and significant, including nearly all pairwise correlations among the limbic structures. Together, these patterns indicate that at deep divergences the limbic structures changed in concert with one another as they covaried inversely with the neocortex.

## Uncertainty quantification with Bayesian inference

Phylogenetic independent contrasts yields a point estimate and p-value for each correlation, but lacks a direct measure of uncertainty. This matters especially where the number of contrasts is small. To quantify this uncertainty, we repeated the analysis in a Bayesian framework, which returns a full probability distribution over the correlation rather than a single value.

We modeled brain core, neocortex, and each target limbic structure jointly under a three-trait multivariate Brownian motion model, in which traits drift randomly along the branches of the phylogenetic tree. Under this model, trait values at the tips of the tree follow a multivariate normal distribution, with mean equal to the trait values at the root of the tree and covariance equal to *R C*, where is the Kronecker product (Methods). Here, *C* is the phylogenetic variance–covariance matrix, which is fixed by the tree and represents how much evolutionary history each pair of species shares. *R* is the evolutionary rate matrix, which is estimated from the data and represents how fast each trait evolves and how strongly pairs of traits covary with one another. Markov chain Monte Carlo was used to sample *R*, and the partial correlation between the neocortex and each limbic structure was computed from the sampled matrices [17].

Fitting *R* to the full tree reproduced the pattern seen in the pooled PICs analysis. Between the neocortex and the olfactory bulb, the posterior mean partial correlation was 0.289, with a 95% credible interval of [ 0.444, 0.123] and a *>* 99.9% posterior probability that the correlation is negative. However, for all the other individual limbic components, the 95% credible intervals included zero (Supplementary Table 1), indicating no reliable inverse relationship with the neocortex. Between the neocortex and the limbic system as a whole, the posterior mean partial correlation was 0.097, with a 95% credible interval of [ −0.251, +0.061] and an 89.1% posterior probability that the correlation is negative. Thus, although the posterior distribution for the composite limbic system was shifted toward negative values, the data did not provide strong posterior support for an inverse relationship with the neocortex.

To restrict the analysis to a subset of divergences based on phylogenetic depth, we next estimated the posterior directly from the contrasts after partitioning them by depth (Methods). Applied to all contrasts, this approach recovers the neocortex-olfactory bulb and neocortex-limbic correlations obtained from the rate matrix above, confirming that the two estimate the same quantity. Restricted to deep divergences (≥ 50 Ma, *n* = 20), the correlation between the neocortex and the limbic system becomes decisively negative, with a posterior mean partial correlation of -0.703, a 95% credible interval of [ 0.847, 0.444]), and a *>* 99.99% posterior probability that the correlation is negative. The shallow subsets showed no significant correlation, with a posterior mean of +0.071 (95% CI [ −0.094, +0.231]). Similarly, at deep divergences, negative correlations were found between the neocortex and each limbic component considered individually (Supplementary Table 2). Thus, despite the limited number of contrasts available at deep branches of the evolutionary tree, the inferred relationships are supported with high posterior probability, confirming the conclusions drawn from the PICs analysis.

## Discussion

We used volumetric data across 157 mammalian species and phylogenetic analyses spanning 100 million years of evolutionary divergence to determine when during mammalian evolution the tradeoff between the neocortex and the limbic system emerged. Cross-species comparisons using raw brain volumes recover a strong negative relationship between the two structures, but this relationship weakens substantially and loses significance once shared ancestry is accounted for. When we resolved the signal by evolutionary depth, however, a different picture emerged. The tradeoff is not absent, but concentrated almost entirely at the deepest divergences of the mammalian phylogeny, where it is strong, extends across every limbic structure, and is robust to the choice of depth threshold. The tradeoff between the neocortex and the limbic system therefore appears to be a signature of the reorganizational events that accompanied the earliest radiations of mammalian lineages, rather than a feature that persisted throughout brain evolution.

The depth at which we detect this signal is notable considering the broader history of mammalian diversification. Fossil and phylogenomic evidence indicate a rapid diversification of mammalian lineages following the end-Cretaceous mass extinction 66 million years ago, with stem representatives of most modern orders appearing within a few million years of that event [18, 19]. The oldest crown fossils of several major groups in our dataset date to this window [20–22], around the 50-million-year threshold we use to define deep divergences. This period is widely regarded as one of the most rapid episodes of ecological and morphological diversification in mammalian history [23]. It was accompanied by pronounced differentiation in body form, locomotion, diet and sensory ecology, as lineages radiated into niches left vacant by non-avian dinosaurs. This likely provided opportunities for corresponding changes in the relative investment in major brain systems. Testing whether these morphological and ecological shifts are linked to the neural reallocation we describe would require reconstructing the ecology and cranial anatomy of ancestral lineages from fossil evidence and ancestral state reconstruction techniques [24–26]. Such data could then be related to the timing and magnitude of the neocortex–limbic tradeoff we report, at the nodes where the tradeoff is strongest.

Our results also speak to a longstanding distinction between concerted and mosaic accounts of brain evolution. Concerted evolution, in its classical interpretation, holds that developmental linkages tie brain structures together such that change in one component entails coordinated change across the brain as a whole [1, 2]. Mosaic evolution instead emphasizes selection acting on interconnected components serving a particular function, largely in isolation from the rest of the brain [4, 5]. Neither model, taken strictly, describes what we observe. Under a strict mosaic account, each limbic structure would evolve independently, shaped by selective pressures specific to its own function. Despite the diversity of limbic structures, we find that these structures covary tightly with one another, and covary negatively with the neocortex at the deepest divergences. A strict whole-brain concerted account, in turn, would imply that the neocortex should scale together with limbic structures as part of a single, shared developmental program. What we observe instead is a hybrid pattern: concerted change within the limbic system, paired with a tradeoff with the neocortex, based broadly on function [3, 7]. This suggests that whatever process links the neocortex and limbic system operates at the level of these two functional systems, rather than the whole brain or individual finer-scale structures. Consistent with this, coupling between limbic system components has been linked to changing sensory demands over vertebrate evolution, including the transition to terrestrial life [27, 28].

The deep-node concentration of the tradeoff may reflect distinct developmental origins of the neocortex and the limbic structures within the early telencephalic pallium. The prosomeric model, which describes forebrain organization in terms of molecularly defined developmental compartments [29, 30], holds that the alar telencephalon subdivides from early neural plate stages to distinct pallial domains, consisting of medial, dorsal, lateral, and ventral sectors. The medial sector gives rise to the hippocampal formation, the dorsal sector to the neocortex, and the lateral and ventral sectors to olfactory and amygdalar derivatives. It has been proposed that differential expansion of these adjacent domains, rather than an ongoing interaction between already-differentiated structures, could be the mechanistic basis of the neocortex-limbic tradeoff [6, 7]. This developmental account is consistent with our observation that the tradeoff is concentrated at the deepest phylogenetic nodes. If the relative allocation of pallial territory to neocortical versus limbic domains is set largely by patterning events early in development [31, 32], divergence in that allocation would be expected to arise chiefly at branch points marking major shifts in developmental programs [33, 34], rather than accumulating gradually within lineages.

This account of shared origin and pallial allocation, however, leaves open the question of why there are separate pallial domains and why they diverge into fundamentally different kinds of circuits. Arguably, the answer lies in a correspondence between statistics and structure. The structure of neocortical circuits is organized around topographic wiring, in which nearest-neighbor relationships within sensory representations are preserved as they project through the thalamus into the cortex and across successive cortical areas [35–40]. The precision of this mapping coarsens progressively from primary sensory areas toward higher-order and frontal cortex, but the organizational principle is maintained. Limbic structures, by contrast, largely lacks this preservation of spatial adjacency, instead utilizing distributed connectivity through which correlated but spatially dispersed inputs are bound [41–46]. These wiring patterns, established early in development, serve as innate structure, or inductive biases [7, 47–49], adapted to the spatial statistics of the information represented in each system. Given their distinct wiring, the two systems cannot co-inhabit the same pallial domain. Allocating capacity to one may therefore come at the expense of the other under constraints on energy and space.

## Methods

### Data

Volumes of brain structures were compiled from published comparative datasets. Data for insectivores, prosimians, and primates were obtained from [11, 12]; macrobats from [13]; carnivores, pinnipeds, ungulates, and xenarthrans from [6]; and *Mus musculus* from [14]. The datasets include volumes of a core set of brain structures—the medulla, mesencephalon, diencephalon, and striatum—as well as the neocortex, cerebellum, limbic system components, and olfactory structures. Olfactory structures comprise the olfactory bulb and the olfactory cortex. The latter includes the primary olfactory (piriform) cortex, the olfactory tubercle, and associated basal forebrain nuclei [6, 12]. Limbic components are the amygdala, septum, hippocampus, and schizocortex. The schizocortex consists of the entorhinal cortex and associated cortical structures (parahippocampal, presubicular, parasubicular, and subicular cortices). We grouped the olfactory structures with the limbic components and refer to the combined set as the limbic system [6, 7]. All brain-region volumes were log-normalized before analysis.

We excluded *Geogale aurita*, an insectivore with an unusually small neocortex [11], as an outlier. Microbats reported in [13] were also excluded because of their comparatively small overall brain size, which may reflect special selection for nimble-bodied flight [6]. These exclusions yielded a dataset of 176 species. Nineteen species lacked divergence-time information in TimeTree and therefore could not be placed on the phylogeny (see Phylogenetic trees), leaving 157 species with both volumetric and phylogenetic data.

Allometric relationships among species values, before phylogenetic correction, were estimated by ordinary least-squares regression on log-transformed volumes, ln(*V*_structure_) = *α* + *β* ln(*V*_core_), where *V*_structure_ is the volume of the brain structure of interest and *V*_core_ is the volume of brain core.

### Phylogenetic trees

Phylogenetic trees with branch lengths proportional to evolutionary time were constructed using PareTT [50], which retrieves pairwise molecular divergence-time estimates from the TimeTree database [10] and assembles them into dated ultrametric trees. Separate trees were constructed for each of the eight taxonomic groups, along with a combined tree containing all species (Figure 2A). The tree was constructed to be ultrametric, such that all terminal nodes were equidistant from the root to within 1% of the mean root-to-tip distance. Node ages in the resulting trees were spot-checked against reference divergence times in TimeTree. The full phylogenetic tree is shown in Supplementary Figure 4.

### Phylogenetically independent contrasts

Species values cannot be treated as independent data points, because closely related species share part of their evolutionary history. Phylogenetically independent contrasts (PICs) remove this dependence by converting the *n* species values at the tips of the tree into *n* − 1 contrasts (comparisons), one at each internal node, that are statistically independent of one another [16, 51].

The method is based on a Brownian motion model of evolution in which a trait (in our case, the log-transformed volume of a brain structure) changes along each branch of the tree by a random increment drawn from a normal distribution with mean zero and variance *σ*^2^*t*, where *t* is the branch length and *σ*^2^ is the rate of evolution. Consider two sister species *i* and *j*, with trait values *x_i_* and *x_j_*, that diverged from a common ancestor, with branch lengths *t_i_* and *t_j_* from that ancestor. Since their divergence, each species has accumulated its own independent random change, with variance *σ*^2^*t_i_* and *σ*^2^*t_j_* respectively. The difference between them, *x_i_ x_j_*, reflects only evolution that occurred after the split. Under Brownian motion, it has mean zero and variance *σ*^2^(*t_i_* + *t_j_*). Pairs separated by longer branches have more time to diverge and therefore are expected to differ more. The typical size of the difference, its standard deviation, is the square root of the variance, and therefore grows as 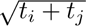. Dividing the observed difference by this quantity gives the standardized contrast

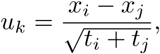

which expresses each difference relative to the difference expected for that amount of divergence. Standardized in this way, the contrasts at all nodes can be treated as independent draws from the same distribution (normal with mean zero and variance *σ*^2^), regardless of branch length.

To compute contrasts at deeper nodes, the value at each internal node is estimated from its two descendants as a weighted average,

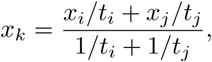

which gives more weight to the descendant on the shorter branch, since it is expected to resemble the ancestor more closely. This estimate deviates from the true ancestral value by a random error. Under Brownian motion, the variance of this error is equivalent to an additional length of *t_i_t_j_/*(*t_i_* + *t_j_*) on the branch leading to node *k*. This branch is therefore lengthened by that amount, and the resulting effective branch length is used when node *k* is compared with its own sister. Repeating these steps from the tips toward the root (a postorder traversal) yields one contrast per internal node, 156 in total for our 157 species. We implemented this procedure in Python using DendroPy [52], and computed contrasts separately for the log-transformed volumes of brain core, neocortex, limbic system, and each limbic component. Each contrast was labeled with the age of the node at which it was computed, which allowed contrasts to be partitioned by phylogenetic depth (see Deep divergence analysis).

To test whether the contrasts were standardized adequately, we correlated the absolute value of each standardized contrast with the amount of change expected from its associated branches [53]. No trait showed a significant relationship (brain core, neocortex, limbic system, and each limbic component; all *p >* 0.29), confirming that the standardization was adequate.

Contrasts of different brain structures at the same node measure correlated evolutionary change along the same branches. When the brain core enlarges along a branch, most structures enlarge with it (Figure 1a). Thus, contrasts of any two structures would correlate positively simply through their shared dependence on brain core. To remove this effect, the contrasts of each structure were regressed against the contrasts of brain core. Correlations between the residuals of two structures measure whether they evolved together, independently of brain core and shared ancestry. All regressions and correlations of contrasts were computed through the origin [53].

### Statistical significance

Statistical significance was assessed using a two-tailed t-test with *df* = *n* − 2 for correlations between contrasts, where *n* is the number of contrasts, and *df* = *n* − 3 for species-level correlations, where *n* is the number of species. Thresholds: \**p <* 0.05, \*\**p <* 0.01, \*\*\**p <* 0.001, ψ 0.05 ≤ p < 0.10.

### Deep divergence analysis

To examine whether the neocortex–limbic tradeoff is concentrated at particular periods of evolutionary history, PICs were partitioned by node age. A primary threshold of 50 Mya was used to divide contrasts into deep (≥50 Mya, *n* = 20 contrasts) and shallow (*<*50 Mya, *n* = 136 contrasts) subsets, and the correlation was computed separately within each (Figure 3). The dependence of this relationship on threshold choice was then assessed by sweeping node-age thresholds from 5 Mya upward in 2-million-year increments (Figure 4). At each threshold, the correlation coefficient was computed using all contrasts with node ages older than that threshold. The sweep was truncated at 63 Mya, beyond which too few contrasts remained for reliable estimation. The 50 Mya primary threshold falls within the range over which the correlation had plateaued (approximately 48 Mya onward), so the deep–shallow comparison is not sensitive to the exact choice of threshold (Supplementary Figure 5).

### Bayesian inference

To complement the frequentist PICs analysis and quantify uncertainty in the evolutionary covariance estimates, we performed a Bayesian Markov chain Monte Carlo (MCMC) analysis under a multivariate Brownian motion model [17]. Starting from the log-transformed volumes of each brain structure at the tips of the tree (the 157 extant species), we estimated the covariances between the structures along the branches of the tree.

Under Brownian motion, a trait changes along each branch by a random increment drawn from a normal distribution with mean zero and variance *σ*^2^*t*, where *t* is the branch length and *σ*^2^ is the evolutionary rate of the trait. At each tip of the tree, the trait value equals the root value plus the sum of the increments along the path from the root to that tip. Because these increments are independent zero-mean normal variables, the set of tip values follows a multivariate normal distribution with mean equal to the root value. Two species share the same increments only along the branches they have in common, and therefore covary in proportion to their shared evolutionary history. The covariance of the tip values is consequently *σ*^2^*C*, where *C* is the 157 × 157 phylogenetic variance–covariance matrix for the 157 species in our dataset. The diagonal entries of *C* are the root-to-tip path lengths, and each off-diagonal entry *C_ij_* is the path length shared by species *i* and *j*, from the root to their most recent common ancestor.

With three traits (brain core, neocortex, and a target limbic structure), the scalar *σ*^2^ is replaced by a 3 × 3 rate matrix *R*. The diagonal entries of *R* are the evolutionary rates of the three traits, and the off-diagonal entries are their evolutionary covariances. Along each branch, the three traits change by a random increment vector drawn from a multivariate normal distribution with mean zero and covariance *Rt*. As in the single-trait case, summing these increments along the branches produces a multivariate normal distribution at the tips, with mean equal to the root trait values and covariance *V* = *R C*, where denotes the Kronecker product.

Each entry of matrix *V* is the product of how strongly two traits covary per unit time (*R_kl_*) and how much evolutionary history two species share (*C_ij_*). *C* is fixed by the time-calibrated tree, but *R* needs to be estimated. The root values of the three traits also need to be estimated. We arrange these root values in a vector *A*. For any candidate *A*, subtracting the root value from the observed tip values of a trait gives the summed evolutionary increments from the root to each tip. The likelihood of these increments for a candidate *R* is given by the multivariate normal density with mean zero and covariance *V*. We estimated *A* and *R* using MCMC, as described below. The partial correlation between the neocortex and the target limbic structure, controlling for brain core, was then calculated from posterior samples of *R*. The procedure was repeated independently for each limbic structure.

For MCMC, parameters in *A* and *R* were sampled using Metropolis-Hastings with single-component updates, initialized at maximum likelihood estimates. Flat priors were placed on the elements of *A* and *R*, and proposals for which *R* was not positive definite were rejected. Chains were run for 100,000 generations with 10,000 burn-in, retaining every 100th sample for 900 posterior samples per chain [54, 55]. Three independent chains (seeds 42, 123, 125) were run per comparison. Convergence was confirmed by Gelman-Rubin *R*^^^ *<* 1.01 [56].

### Posteriors by node age

The rate matrix *R* described above is fitted to the tree as a whole, and so yields a single evolutionary correlation with no dependence on node age. To assess whether the correlation differs between deep and shallow divergences, we estimated the posterior from the PICs instead. PICs are sufficient statistics for the same Brownian motion model and each contrast carries the age of the node at which it was computed. We partition the contrasts into deep (≥ 50 Mya) and shallow (*<* 50 Mya) subsets.

Within each subset, the brain-core-residualized contrasts of the neocortex and of the target structure were treated as draws from a bivariate normal distribution, and we estimated the correlation *ρ* between them. This leaves a single unknown parameter, so the posterior can be obtained exactly rather than by sampling. For this, we placed a uniform prior on *ρ* over ( −1, 1), evaluated the posterior density on a grid of 6,000 values spanning that interval, and normalized it by numerical integration. We report the posterior mean, the 95% credible interval, and the posterior probability that *ρ <* 0.

## Supplementary Materials

The Supplementary Materials contain eight figures (Figures S1-S8) and two tables (Tables S1 and S2) that are referenced in the main text and presented below.

**Supplementary Figure 1:**
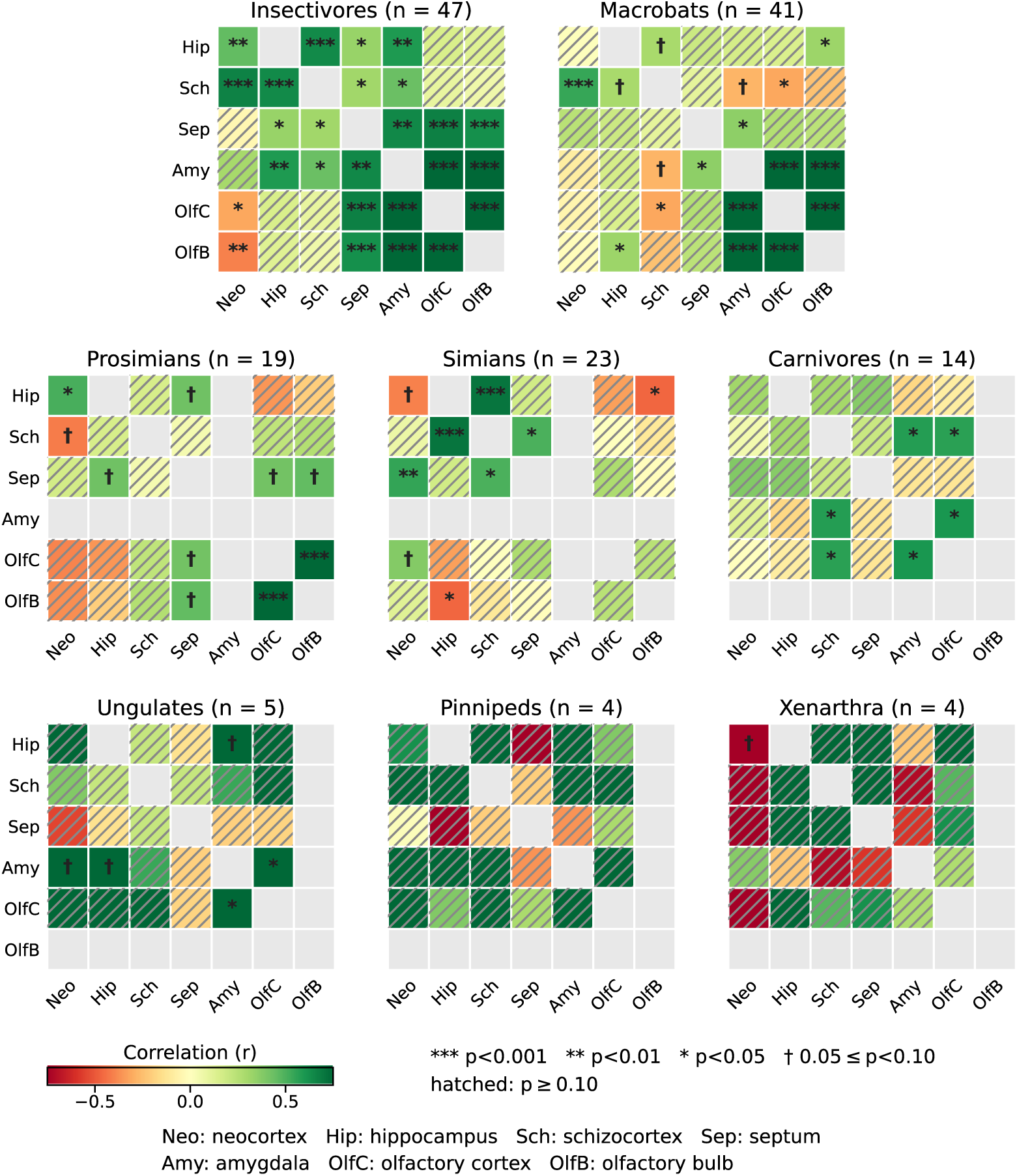
Covariation among brain structures within individual taxonomic groups. Each panel represents a single taxon. Each cell within a panel shows the residual correlation between two brain structures after regressing their volumes against the volume of brain core. The number of species in each taxon is given in brackets. Color indicates correlation coefficient. Gray cells mark a structure against itself or a structure not measured separately in that taxon.

**Supplementary Figure 2:**
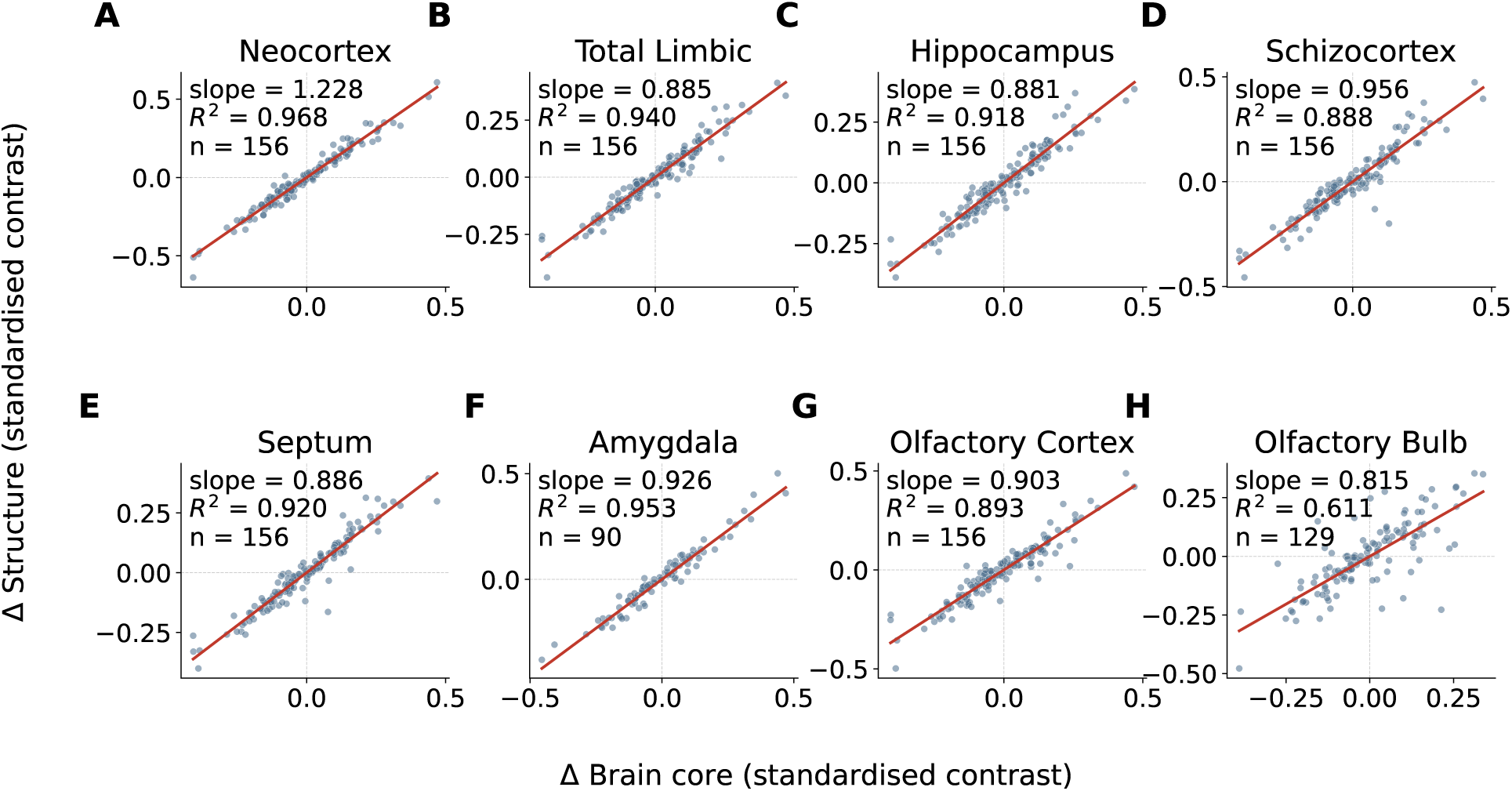
Regression of each structure’s contrasts against brain-core contrasts, through the origin. Standardized contrasts for each structure (vertical axis) against standardized contrasts of brain core (horizontal axis). The regression line (red) is fit through the origin. The residuals from these fits represent brain structure volumes that account for both allometric scaling and phylogenetic non-independence. The number of contrasts differs between panels because amygdala and olfactory bulb are measured in fewer species in our dataset. The neocortex is the only structure with a slope above one (1.228), meaning it grows faster than brain core across the tree, while every limbic structure scales more slowly (0.815-0.956). The olfactory bulb shows the weakest coupling to brain core (*R*^2^ = 0.611, against 0.888-0.968 for all other structures).

**Supplementary Figure 3:**
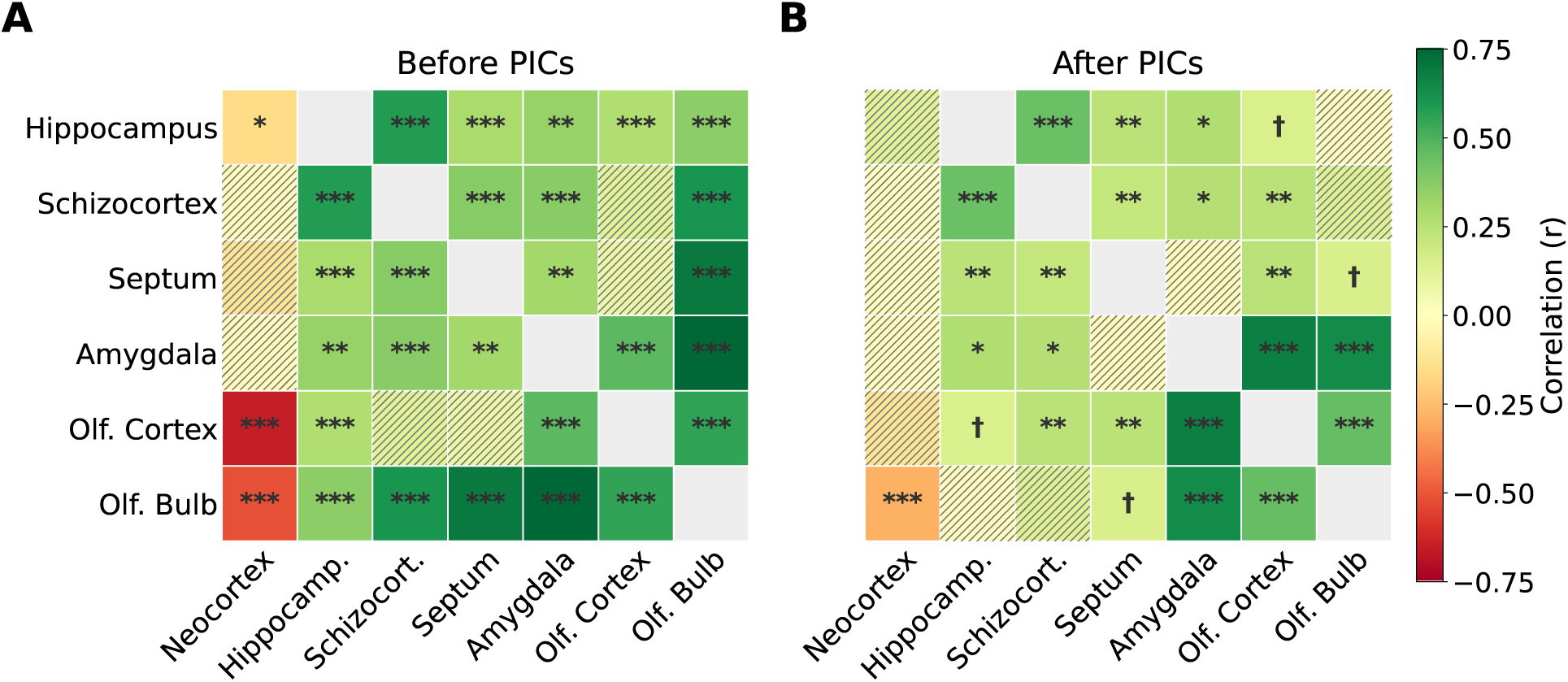
Pairwise correlations among all structures, before and after phylogenetic correction. (A) Correlations between brain structure volumes across the 157 species. The volume of each structure was residualized on brain-core volume by ordinary least-squares regression with an intercept. The residuals were then correlated by Pearson’s *r*. These are species values rather than contrasts. (B) The same pairs after phylogenetic correction. Contrasts were residualized on brain-core contrasts by regression through the origin. The residuals were then correlated through the origin [53]. Layout, ordering and color scale match Figure 1c. Cells are hatched where *p ≥* 0.10. Gray cells are the structure against itself. Before phylogenetic correction, the neocortex covaries negatively with the olfactory bulb, olfactory cortex and hippocampus. After correction, only the neocortex–olfactory bulb relationship remains significant. Five of the fifteen limbic–limbic pairs are also significant before correction but not after: hippocampus– olfactory bulb, septum–olfactory bulb, schizocortex–olfactory bulb, septum–amygdala, and hippocampus–olfactory cortex. ^∗^*p <* 0.05, ^∗∗^*p <* 0.01, ^∗∗∗^*p <* 0.001, **^ψ^**0.05 *≤ p <* 0.10.

**Supplementary Figure 4:**
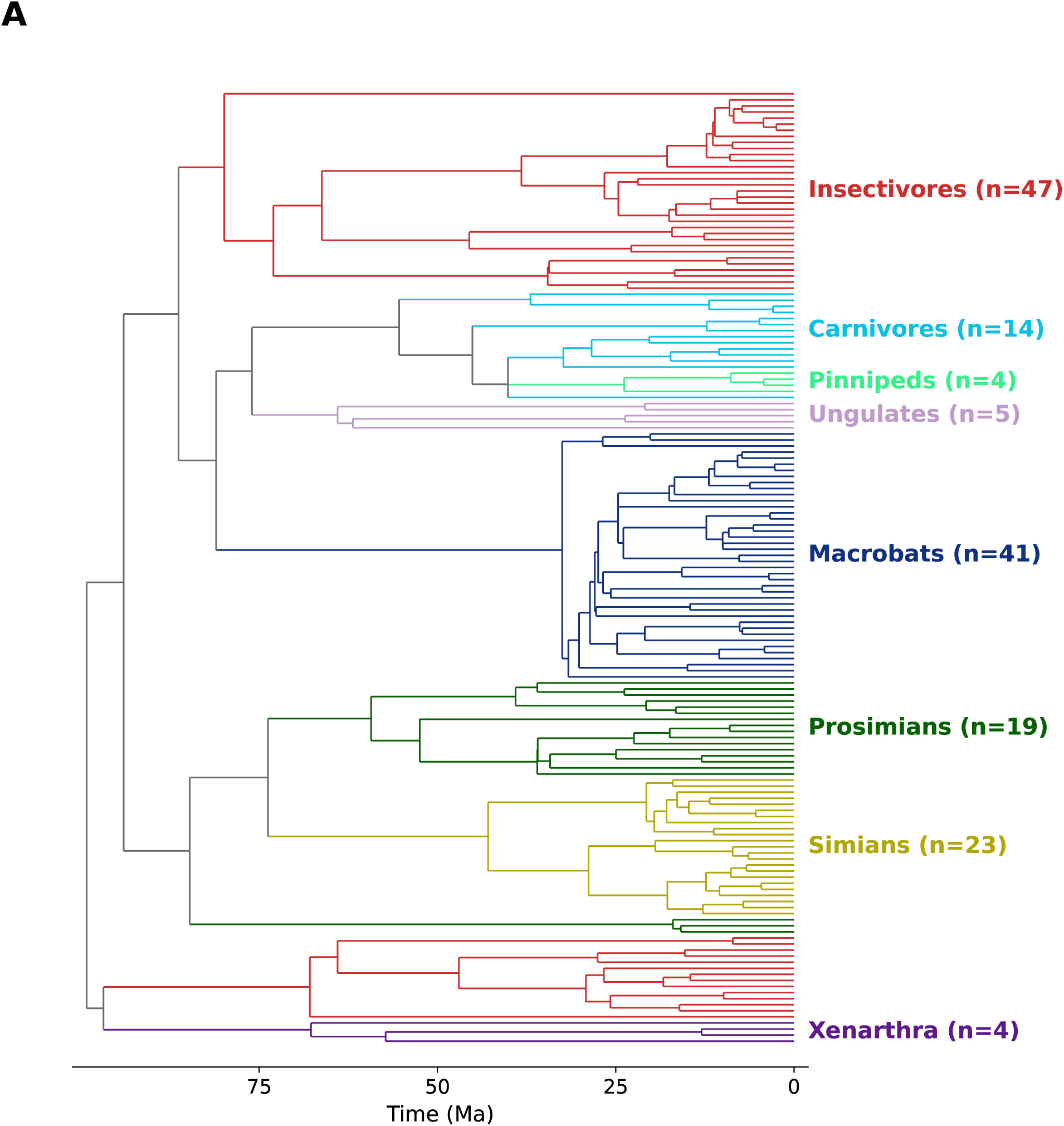
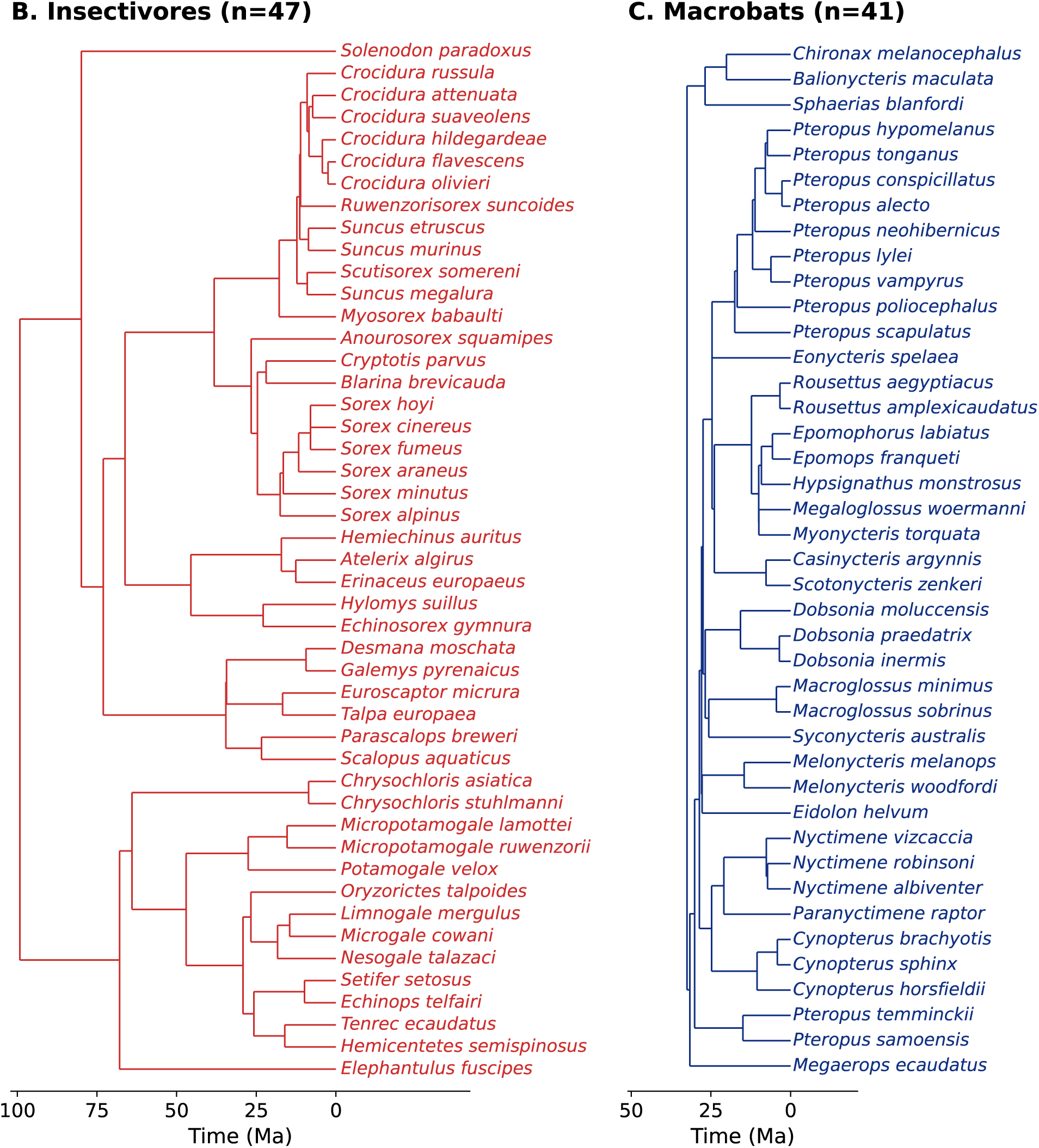
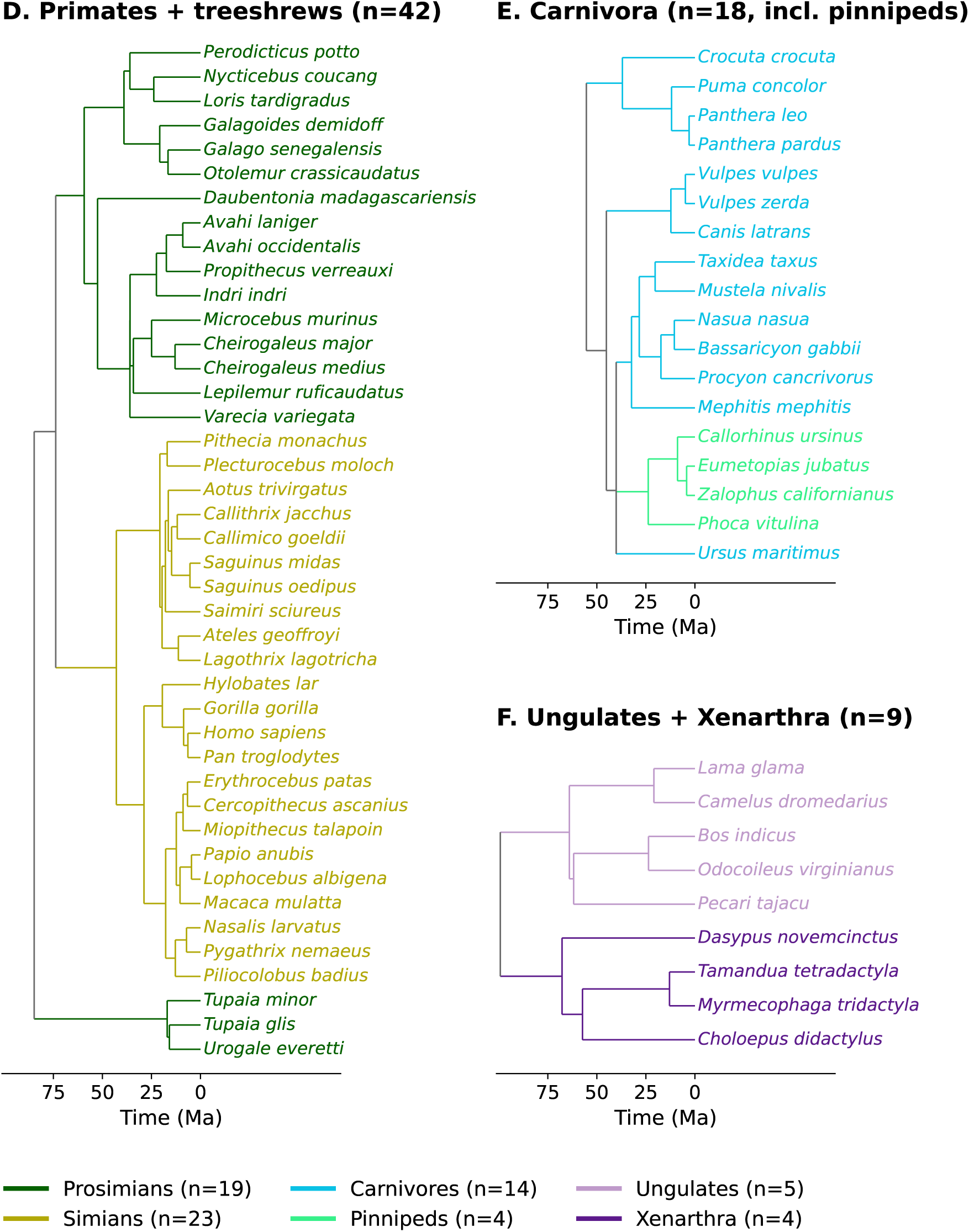
Time-calibrated phylogeny of the 157 species in the dataset. (A) The full tree, with branches colored by taxonomic group. Tip labels are omitted in this subplot. Branches whose descendants span more than one group are drawn in gray. Branch lengths are divergence times in millions of years, obtained from PareTT and cross-checked against TimeTree. The root is at 99.2 Mya. Clades run top to bottom in the same order as the condensed tree in Figure 2A. Three groups are not contiguous in the tree—insectivores, prosimians and carnivores. This is the full tree underlying every contrast computed in this study. The figure is continued in the next two pages. The same tree broken out by group, with species labels: (B) insectivores (*n* = 47), (C) macrobats (41). The Panels are drawn at a common scale, so divergence times are directly comparable between panels and all tips align at the present. Each axis extends back to the root of the clade shown. (D) Primates and treeshrews (42), (E) Carnivora, including the pinnipeds nested within it (18), (F) ungulates and xenarthrans (9). The Panels are drawn at a common scale, so divergence times are directly comparable between panels and all tips align at the present. Taxonomic groups follow the sheet-level grouping of the source dataset, so the prosimian group includes three treeshrews (*Tupaia minor*, *Tupaia glis* and *Urogale everetti*), which are Scandentia rather than strepsirrhine primates. These appear at the bottom of (D).

**Supplementary Figure 5:**
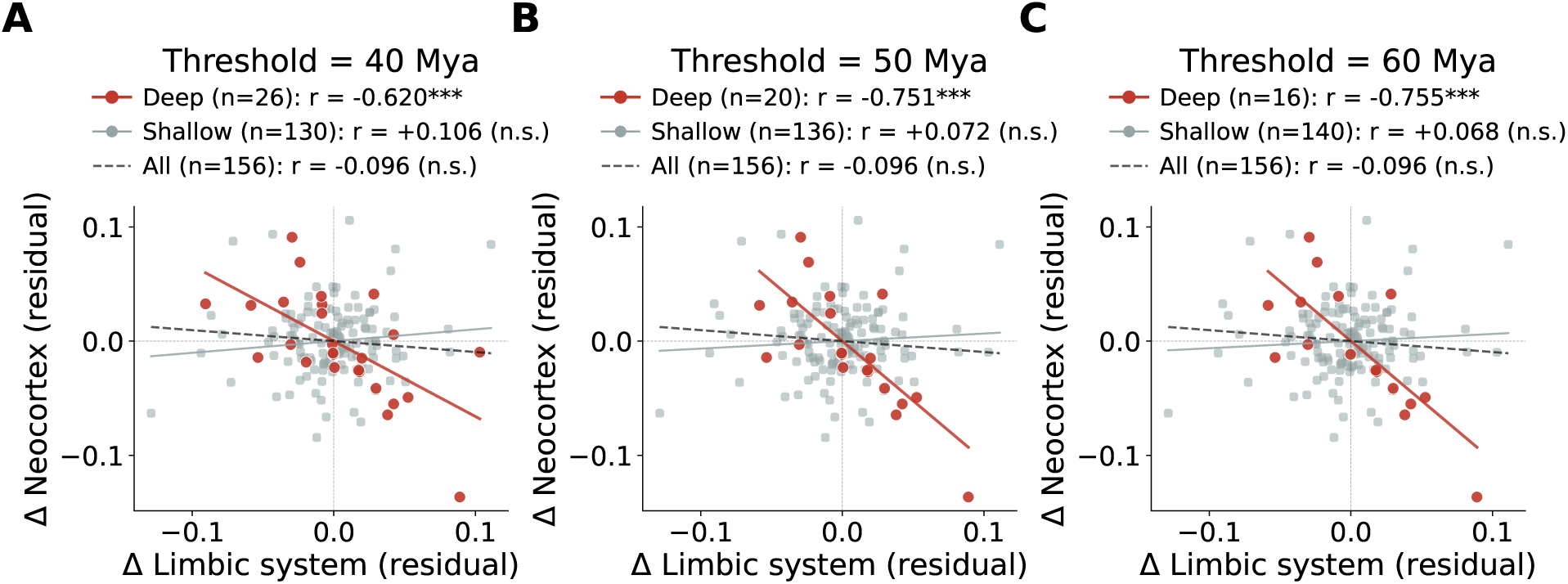
The deep/shallow partition is robust to the choice of threshold. Neocortex against total limbic system, in contrast residual space, with contrasts split into deep and shallow subsets at (A) 40 Mya, (B) 50 Mya and (C) 60 Mya. Red points and line show deep contrasts. Gray points and line show shallow contrasts. Lines are fit through the origin. The correlations are also calculated through the origin (i.e. without subtracting the means of the variables) [53]. The deep subset is strongly and significantly negative at every threshold (*r* = *−*0.620, *−*0.751 and *−*0.755; all *p <* 0.001), whereas the shallow subset is non-significant at every threshold (*r* = +0.106, +0.072 and +0.068; all *p >* 0.22). Panel B reproduces Figure 3 of the main text. ^∗^*p <* 0.05, ^∗∗^*p <* 0.01, ^∗∗∗^*p <* 0.001, **^ψ^**0.05 *≤ p <* 0.10.

**Supplementary Figure 6:**
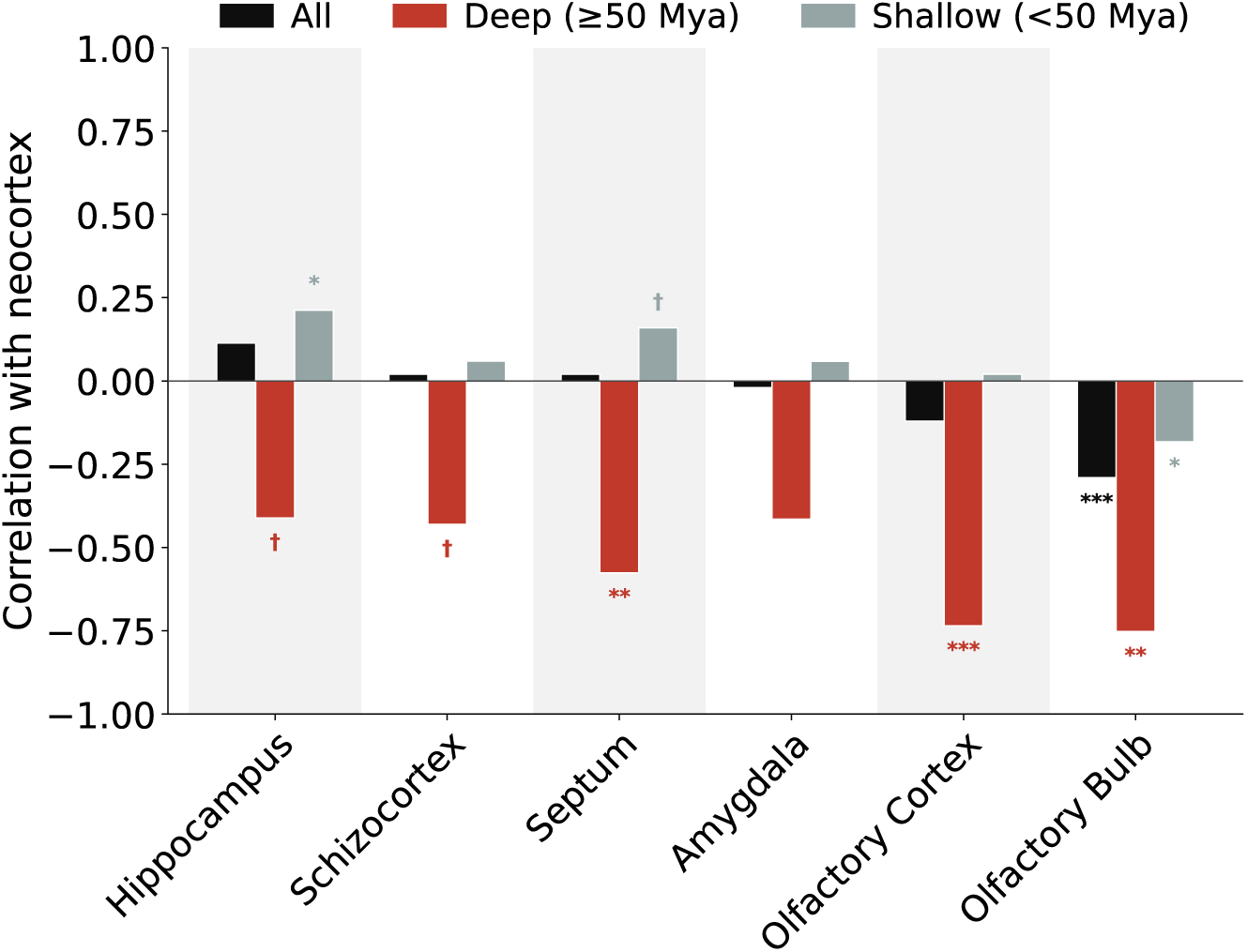
Neocortex against each limbic component at deep and shallow nodes. Shown are correlations between neocortex contrasts and the contrasts of each limbic structure, computed through the origin, after residualizing both on the contrasts of brain core. Black, all 156 contrasts; red, deep contrasts (*≥* 50 Mya); gray, shallow contrasts (*<* 50 Mya). The deep subset contains 20 contrasts for hippocampus, schizocortex, septum, and olfactory cortex, and 12 contrasts for amygdala and olfactory bulb. The number of contrasts differs because the amygdala and olfactory bulb are not measured separately in all the species in the source dataset. At deep nodes every component covaries negatively with the neocortex. Bars whose absolute value is below 0.02 are drawn at 0.02 so that they remain visible. ^∗^*p <* 0.05, ^∗∗^*p <* 0.01, ^∗∗∗^*p <* 0.001, **^ψ^**0.05 *≤ p <* 0.10.

**Supplementary Figure 7:**
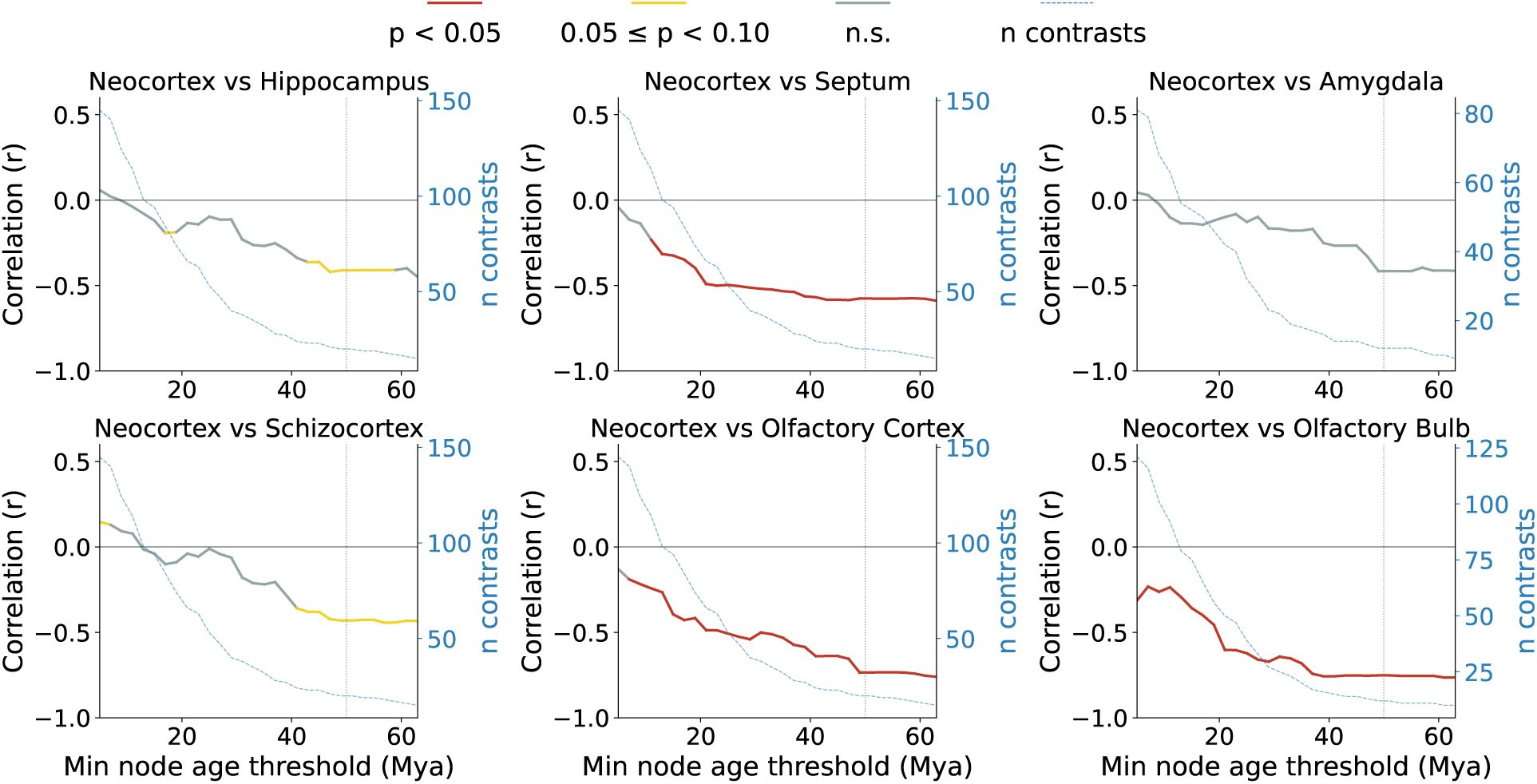
Neocortex–limbic correlations across increasing evolutionary depth. Correlations are computed from PICs, residualized on brain-core to remove the effect of brain size (see main text). Plotted are correlations (left axis, solid line) as a function of minimum node-age threshold (bottom axis). At each threshold, only contrasts at nodes older than the threshold are included. The number of contrasts remaining at each threshold is also shown (right axis, blue dashed). The dotted vertical line marks the 50 Mya threshold used in the deep vs shallow analyses. The olfactory bulb and olfactory cortex, with the strongest deep-node correlations (r=-0.75 and -0.74, respectively), were significant across a broad range of thresholds, as was the septum. The hippocampus and schizocortex, whose deep-node effects were genuinely weaker (r*≈*-0.42), reached significance within a narrower window (approximately 46–54 Ma). At shallower thresholds their correlation was diluted by the many shallow contrasts, and at deeper thresholds too few contrasts remained to detect an effect of that magnitude. The weaker significance of these structures thus reflects smaller effect sizes and limited statistical power at the deepest nodes.

**Supplementary Figure 8:**
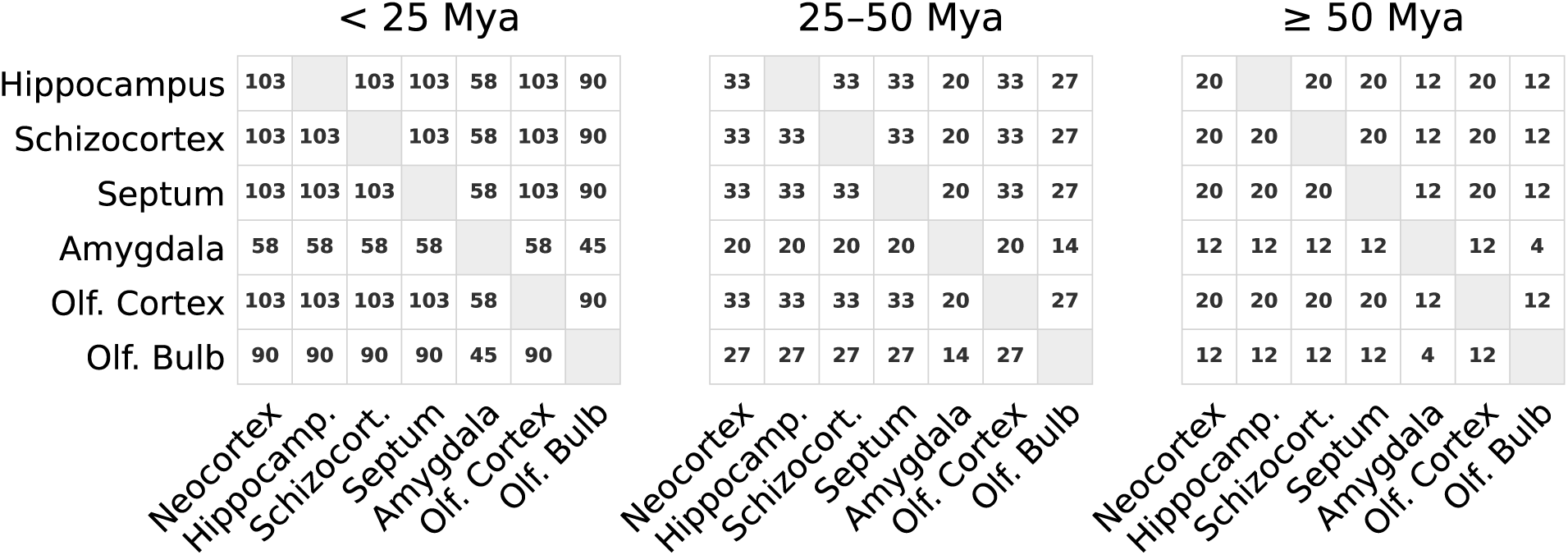
Number of contrasts used for each pairwise correlation in Figure 4C of the main text. Layout and node ranges match Figure 4C. Each cell gives the number of contrasts available for that pair of structures. This number depends on the number of nodes at the specified phylogenetic depths and the availability of measurements for both structures at the tips of the tree. Pairs involving the amygdala or olfactory bulb have fewer contrasts because these structures were measured separately in fewer species. At *≥* 50 Mya, most pairs have 20 contrasts, pairs involving the amygdala or olfactory bulb have 12, and the amygdala–olfactory bulb pair have 4.

**Supplementary Table 1:** Posterior partial correlations between the neocortex and each limbic structure, controlling for brain core. Estimated from a three-trait evolutionary rate matrix *R* fitted to the full tree by MCMC (100,000 generations, 10,000 burn-in, sampled every 100, three chains with seeds 42, 123 and 125; Methods). “% neg” is the proportion of pooled posterior samples below zero. *R*^^^ is the Gelman–Rubin potential scale reduction factor; values below 1.01 indicate convergence. The olfactory bulb is the only structure whose 95% credible interval excludes zero.

| Structure | $n$ | Posterior mean | 95% credible interval | % neg | $\hat{R}$ |
| --- | --- | --- | --- | --- | --- |
| Total Limbic | 157 | −0.097 | [−0.251, +0.061] | 89.1% | 1.000 |
| Hippocampus | 157 | +0.115 | [−0.043, +0.270] | 7.8% | 1.001 |
| Septum | 157 | +0.014 | [−0.152, +0.173] | 43.3% | 0.999 |
| Amygdala | 91 | −0.009 | [−0.213, +0.196] | 53.7% | 1.000 |
| Schizocortex | 157 | +0.002 | [−0.158, +0.165] | 48.8% | 1.000 |
| Olfactory Bulb | 130 | −0.289*** | [−0.444, −0.123] | 99.96% | 1.000 |
| Olfactory Cortex | 157 | −0.118 | [−0.271, +0.036] | 93.1% | 1.000 |

\*\*\* credible interval excludes zero with > 99.9% of samples on one side.

**Supplementary Table 2:** Posterior correlations between the neocortex and each limbic structure at deep and shallow evolutionary depths. Estimated from the brain-core-residualised contrasts after partitioning them 50 Mya. The contrasts in each subset are treated as draws from a bivariate normal and the posterior for the correlation obtained exactly on a grid under a uniform prior (Methods). “% neg” is the posterior probability that the correlation is negative. Every structure is negative at deep nodes. The number of contrasts differs between structures because amygdala and olfactory bulb are measured in fewer species.

| Structure | Subset | $n$ | Posterior mean | 95% credible interval | % neg |
| --- | --- | --- | --- | --- | --- |
| Total limbic | Deep ($\geq 50$ Mya) | 20 | -0.697 | [-0.843, -0.440] | 100.0% |
| | Shallow ($< 50$ Mya) | 136 | +0.070 | [-0.094, +0.228] | 20.2% |
| Olfactory bulb | Deep ($\geq 50$ Mya) | 12 | -0.658 | [-0.856, -0.270] | 99.8% |
| | Shallow ($< 50$ Mya) | 117 | -0.175 | [-0.335, -0.000] | 97.5% |
| Olfactory cortex | Deep ($\geq 50$ Mya) | 20 | -0.678 | [-0.832, -0.409] | 100.0% |
| | Shallow ($< 50$ Mya) | 136 | +0.008 | [-0.155, +0.170] | 46.4% |
| Hippocampus | Deep ($\geq 50$ Mya) | 20 | -0.348 | [-0.630, +0.033] | 96.4% |
| | Shallow ($< 50$ Mya) | 136 | +0.205 | [+0.044, +0.352] | 0.7% |
| Septum | Deep ($\geq 50$ Mya) | 20 | -0.507 | [-0.733, -0.158] | 99.6% |
| | Shallow ($< 50$ Mya) | 136 | +0.153 | [-0.010, +0.305] | 3.2% |
| Schizocortex | Deep ($\geq 50$ Mya) | 20 | -0.365 | [-0.642, +0.013] | 97.1% |
| | Shallow ($< 50$ Mya) | 136 | +0.057 | [-0.106, +0.217] | 24.6% |
| Amygdala | Deep ($\geq 50$ Mya) | 12 | -0.322 | [-0.667, +0.150] | 91.5% |
| | Shallow ($< 50$ Mya) | 78 | +0.055 | [-0.157, +0.260] | 30.5% |

